# Curvature domains in V4 of Macaque Monkey

**DOI:** 10.1101/2020.02.28.969329

**Authors:** Jiaming Hu, Xue Mei Song, Qiannan Wang, Anna Wang Roe

## Abstract

An important aspect of visual object recognition is the ability to perceive object shape. How the brain encodes fundamental aspects of shape information remains poorly understood. Models of object shape representation describe a multi-stage process that includes encoding of contour orientation and curvature. While modules encoding contour orientation are well established (orientation domains in V1 and V2 visual cortical areas), whether there are modules for curvature is unknown. In this study, we identify a module for curvature representation in area V4 of monkey visual cortex and illustrate a systematic representation of low to high curvature and of curvature orientation, indicative of curvature hypercolumns in V4. We suggest that identifying systematic modular organizations at each stage of the visual cortical hierarchy signifies the key computations performed.

**Significance:** We use intrinsic signal optical imaging in area V4 of anesthetized macaque monkey to study the functional organization of curvature representation. We find a modular basis for cue-invariant curvature representation in area V4 of monkey visual cortex and illustrate a systematic representation from low to high curvature and of curvature orientation, replete with curvature pinwheels. This is the first report of systematic functional organization for curvature representation in the visual system. The use of optical imaging has revealed at a population level spatial details of cortical responses, something which has not been evident from previous studies of single neurons. These data support a representational architecture underlying a ‘curvature hypercolumn’ in V4.

## Introduction

Recognizing the shapes of objects requires information about local contour features, such as orientation and curvature. The encoding of contour orientation in the visual system begins in primary visual cortex (V1) with neurons selective for the orientation of visual contours, and, in the second visual area (V2), with neurons selective for cue-invariant contour orientation (that is, independent of whether contours are defined by luminance, motion, color, or depth cues). These two functions are first embodied in the ‘*orientation domain*’ in V1 (1-3) and the *‘higher order orientation domain’* in V2 (4-5). These signature domains in V1 and V2 mark initial computational elements of shape representation.

We investigate whether there are functional domains representing contour curvature in area V4 of macaque monkey visual cortex, an important intermediate stage of shape encoding. Although both V2 (6-8) and V4 are considered intermediate stages of shape representation, a greater body of neural (9-11) and computational modeling (12-14) evidence points to area V4 as a locus of curvature information processing. V4 neurons are responsive to contour curvature and can be selective for degree of curvature and orientation of curvature (9, 11). However, it is unknown whether such neuronal responses are organized in any way akin to orientation maps in V1 and V2. Previous studies have shown the functional organization of V4 comprises alternating bands of ‘orientation’ and ‘color’ preference (15-17), as well as organization for disparity defined orientation (18), motion direction (16), and spatial frequency (19), but maps for curvature organization have not been demonstrated. Here, we hypothesized that orientation bands in V4 are regions of shape representation that include both orientation and curvature domains.

We used intrinsic signal optical imaging, a method that is well-suited for studying functional domain responses, to examine curvature response in monkey V4. We hypothesized that curvature domains should meet the following criteria: (1) *Distinct response preference*: We expected that curvature domains should prefer curved stimuli over straight stimuli, and, furthermore, would exhibit selectivity for different degrees of curvature (from low to high) or different orientations of a curve. (2) *Curvature maps*: Just as orientation and color domains have distinct maps, we predicted that curvature domains should have distinct spatial distribution. (3) *Cue invariance*: We also predicted that curvature domains would exhibit higher order response properties consistent with the global percept; that is, they should exhibit some degree of cue-invariant curvature response.

## Results

### Curvature domains exist

Using intrinsic optical imaging, we imaged V1, V2, and V4 in three hemispheres of two anesthetized macaque monkeys. As shown in Table S1, almost all experiments described in this study were conducted in at least two cases. Consistent with previous studies (15-17), alternating regions (bands) of color preference vs. orientation preference were observed, with small regions of overlap (Fig. S1). To precisely place our visual stimuli, we also mapped the retinotopy of V4 cortex using 0.2° horizontal and vertical lines and placed the center of visual stimuli (4° in size) on the monitor (Fig. S2, see Methods). We then zoomed in and obtained functional maps of V4. We used oriented achromatic and isoluminant color gratings to obtain maps for orientation, color, and high vs. low spatial frequency (Fig. S3). For all maps, the locations of functional domains were determined by t-value maps (t-map, two-tailed t test, p<0.01) which were calculated by comparing, pixel by pixel, the responses between two different conditions.

We then examined response in V4 to curved vs. straight stimuli, consisting of moving curved (20) or straight sinusoidal gratings (4 straight orientations: 0°, 45°, 90°, 135°; 4 curved orientations: curved up, curved down, curved left, curved right), and flashed single curved (curved up, curved down) or straight lines (horizontal, vertical) (Fig. 1C). To examine preference for curvature over straight gratings, as shown in Fig. 1E, we subtracted curvature (sum of all 4 curvature grating maps) minus straight (sum of all 4 straight grating maps), revealing dark (curvature preferring) domains (two-tailed t test, p<0.01). These curvature maps were distinct from (straight) orientation maps (Fig. 1D, overlay of curvature and straight shown in Fig. 1I). For simplicity of terminology, we will refer to these curvature vs. straight preferring domains as ‘curvature domains’.

**Fig. 1.**
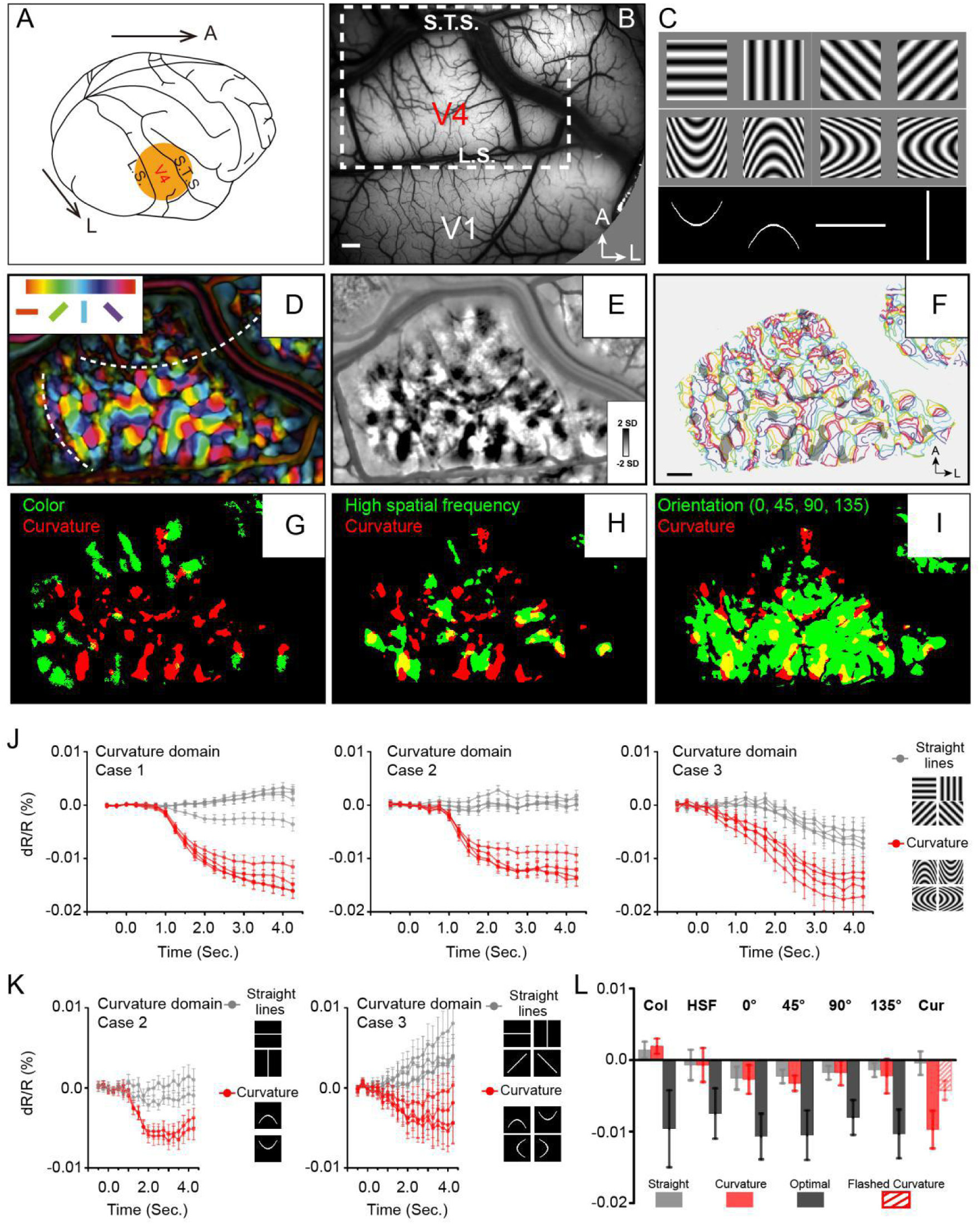
Curvature domains exist and are distinct. A. Location of imaged V4 region (orange disk). L.S., lunate sulcus. S.T.S., superior temporal sulcus, A, anterior, L, lateral. B. View of cortical surface in Case 1. Dotted box: region shown in D-I. C. Stimuli were 4 º in size (see Methods, Fig. S3). D-I: Case 1. D. Color-coded orientation preference map. White dashed lines: approximate borders between color and orientation bands. E. Curvature map: all curved minus all straight gratings. F. Curvature domains (gray patches, two-tailed t-test, p<0.01) superimposed on iso-orientation contours (colored lines, from Fig. 1F). G-I. Overlay of curvature domains (red) and G: color domains (green, from Fig. S3D), H: high spatial frequency domains (green, from Fig. S3E), I: all orientation domains (green, from Fig. S3B, C). J. Response time courses of curvature domains from Case 1 (left), Case 2 (middle), and Case 3 (right). Red lines: preferred stimuli. Gray lines: non-preferred stimuli. K. Response time courses of curvature domains to flashed curved lines. Red timecourses: flashed curved lines. Gray timecourses: flashed straight lines. L. Summary of response amplitudes for color (Col), high spatial frequency (HSF), orientation (0 º, 45º, 90º, 135º), and curvature (Cur) domains shown in J, K. Gray: straight grating. Red: Curved grating. Black: optimal stimulus responses (except for Cur). For Cur, optimal response was to curved gratings (red) and to flashed curved lines (hatched red). Scale bar: 1mm. Error bars: SEM (timecourses in J, K), SD (histogram in L).

If these domains are indeed curvature domains, we predicted that they should be located within the ‘orientation bands’ and should be spatially distinct from straight orientation maps. We found that curvature domains (Fig. 1G-I, red pixels) were distinct from color domains (Fig. 1G, green pixels); they were also distinct from high spatial frequency domains (Fig. 1H, green pixels), suggesting that these curvature responses are not simply due to the high spatial frequency components of the curved grating. In addition, as predicted, the curvature domains fell largely within the orientation band and were in close proximity to the straight orientation domains (Fig. 1I, green pixels). As shown by the yellow pixels (locations of overlap), there is limited overlap between color and curvature domains (Fig. 1G, Case 1, 4.9% of curvature domain locates in color domain; all three cases, 5.7%) and between high spatial frequency and curvature domains (Fig. 1H, Case 1, 26.1% of curvature domain locates in HSF domain; all three cases, 13.9%). However, there is much greater overlap between curvature and orientation domains (Fig.1I, Case 1, 54.9% of curvature domain located in orientation domain; all three cases, 43.0%), raising the possibility that curvature and orientation domains are different components of a shape information processing architecture. When overlaid on outlines of orientation domains, curvature domains appear distinct from orientation domains, and have a semi-regular distribution within the orientation band (Fig. 1F, color: iso-orientation contours, gray patches: curvature domains). Similar curvature vs. straight preference maps were obtained in the other two cases (Figs. S5 and S6).

Note that these curved grating stimuli were generated by periodically repeating an elliptical contour (see Fig. S4); this resulted, for high curvature gratings, high spatial frequency content towards the edge of the stimulus. If the curvature response was primarily due to high spatial frequency, then the curvature map and the high spatial frequency preference should have high overlap. However, this was not observed, suggesting it is unlikely that the curvature response is primarily due to high spatial frequency content (Fig. 1H). We also examined the difference between high curvature degree and low curvature degree gratings, which have different spatial frequency content, are highly overlapped (Fig. S7), suggesting that curvature domain response is not due to spatial frequency content. We also observed that maps were stable over time, indicating these maps are unlikely to be artifactual (Fig. S5). Furthermore, examination of images obtained in response to scrambled curved gratings minus straight gratings did not result instructured maps (Fig. S6), suggesting that, when compared with straight gratings, it is not the presence of multiple orientations or multiple spatial frequencies alone that produced structured maps. In addition, the sizes of curvature domains (mean = 434 μm for all 3 cases, see Fig. S8) fall within the 200-500 μm range of functional domain sizes within V4 (15-17). These data thus suggest that curvature maps are not artifactual and are distinct from previously described functional maps in V4.

We further examined selectivity of response of curvature domains by comparing magnitude of reflectance change in response to curved vs. straight grating stimuli. Timecourses were typical of cortical intrinsic signals in V4, characterized by 2-3 second peak times and 0.01 - 0.03% amplitudes. As expected, color domains exhibited robust response to isoluminant color gratings but weak response to achromatic gratings (Case 1, Fig. S3G, red lines). Similarly, high spatial frequency preference domains responded strongly to achromatic gratings of high spatial frequency but poorly to low spatial frequency gratings (Case 1, Fig. S3H, red lines). Neither color domains nor high spatial frequency domains distinguish curved vs. straight gratings (Wilcoxon test, color domains, p=0.06; high spatial frequency domains, p=0.77, all three cases, Fig.1L). These analyses indicate that curvature processing is not subserved by color or high spatial frequency domains.

We found that preferences of orientation domains and curvature domains were distinct. As expected, straight orientation domains exhibited strongest response to gratings of their respective optimal orientations, relative suppression to gratings of orthogonal orientation, and weak responses to other straight orientations (Fig. S3I-L). In comparison, they displayed relatively weak response to curved gratings (Fig. S3I-L, black lines).

Curvature domains, in contrast, exhibited strong preference for curvature stimuli. This is illustrated in 3 cases of V4 imaging (Fig. 1J). As shown in Fig. 1J left, timecourses of curvature domains revealed strongest amplitudes for curvature gratings (red lines, Case 1, same as case shown in Fig. S3G-L), but relatively weak response to straight gratings (gray lines). Similar preferences are evident for both Case 2 (middle) and Case 3 (right).

To further test whether this differential response is due to curvature, in two cases, we examined response to flashed curved and straight lines. As shown in Fig. 1K, using the same pixels from which significant curved grating responses were obtained, we found that response of curvature domains to single flashed curved lines (red lines), although weaker than that to curved gratings (compare curvature response amplitudes in J and K), was significantly greater than that to single straight lines (gray lines). In contrast, for straight orientation domains (Case 2: Fig. S9E-F, Case 3: Fig. S9H-K) and high spatial frequency domains (Case 2: Fig. S9G, Case 3: Fig. S9L), their responses to flashed curved lines (black, gray) are weak compared to flashed straight lines (orange, blue) (summarized in Fig. S9M, Table S2). Thus, preference for curved over straight contours was demonstrated for both grating and line stimuli, and supports a cue-invariant curvature response.

These results are quantitatively summarized in Fig. 1L. Each of the color domains (Col), high spatial frequency domains (HSF), and orientation domains (0°, 45°, 90°, 135°) exhibit preference for their respective optimal stimulus (black bars), one which far exceeds that to non-preferred stimuli, including that to curved gratings (red bars: curved gratings, Wilcoxon test, p<0.0001). Likewise, curvature domains (Cur) exhibit strong preference for curved gratings (large red bar) and curved lines (hatched red bar) and little response to straight gratings (gray bar). In sum, we find that (1) non-curvature domains are weakly responsive to curvature stimuli and (2) curvature domains are selective for curvature stimuli, to a degree that parallels stimulus preference of other functional domains in V4. This supports the concept that curvature domains in V4 satisfy the ‘selectivity’ criterion for functional domains.

### Mapping types of curvature

#### Low to high curvature maps

To examine whether there could be a continum of curvature domains representing low to high curvatures, we designed a series of curved gratings from low to high curvature (Fig. 2A, upwards: 1-6; downwards: 7-12, Case 1; cf. 20) and presented these stimuli in two cases. Maps obtained in response to these curvature stimuli exhibited regular structure and domain size consistent with functional maps (see Fig. 2A). To examine the possibility of a low to high curvature gradient, we conducted cross correlations between pairs of these images (Fig. 2B, see Methods). We reasoned that, if a gradient of curvature degree exists, then there should be a gradually changing map similarity as curvature degree changes. Correlation values ranged from 0 (no correlation or minor negative correlation, blue) to 1 (two maps are identical, red) (see color bar) (to give reader an idea of what high map correlations values are, the range of correlation values for 4 straight orientation maps in V4 shown in Fig. S10). As shown in Fig. 2B, higher correlation values tended to occur between curvature maps of similar curvature degree (orange and yellow squares in upper left quadrant and lower right quadrant) and lower correlation values between those of dissimilar curvature degree (green and blue squares in upper left quadrant and lower right quadrant). Within this correlation matrix, the lowest correlation values were obtained between opposing curvatures (upper right quadrant and lower left quadrant) and between curved and straight stimuli (rightmost column). High correlation values were obtained between adjacent 1 or 2 curvatures (orange and yellow boxes) (e.g compare values between pairs 2-3, 3-4), but correlation values dropped off with increasing differences in degree of curvature (green and blue boxes) (e.g. 2-3, 2-4, 2-5). This relationship is plotted in Fig. 2C for each of the 6 Up and 6 Down curvature degrees (e.g. Fig. 2C top graph red circles: show declining correlation with curvature difference, see Table S3 for regression values). This declining correlation with curvature difference suggests the presence of a shifting map of curvature degree. Similar results were obtained in a second case (Case 2, Fig. S11A-B).

**Fig. 2.**
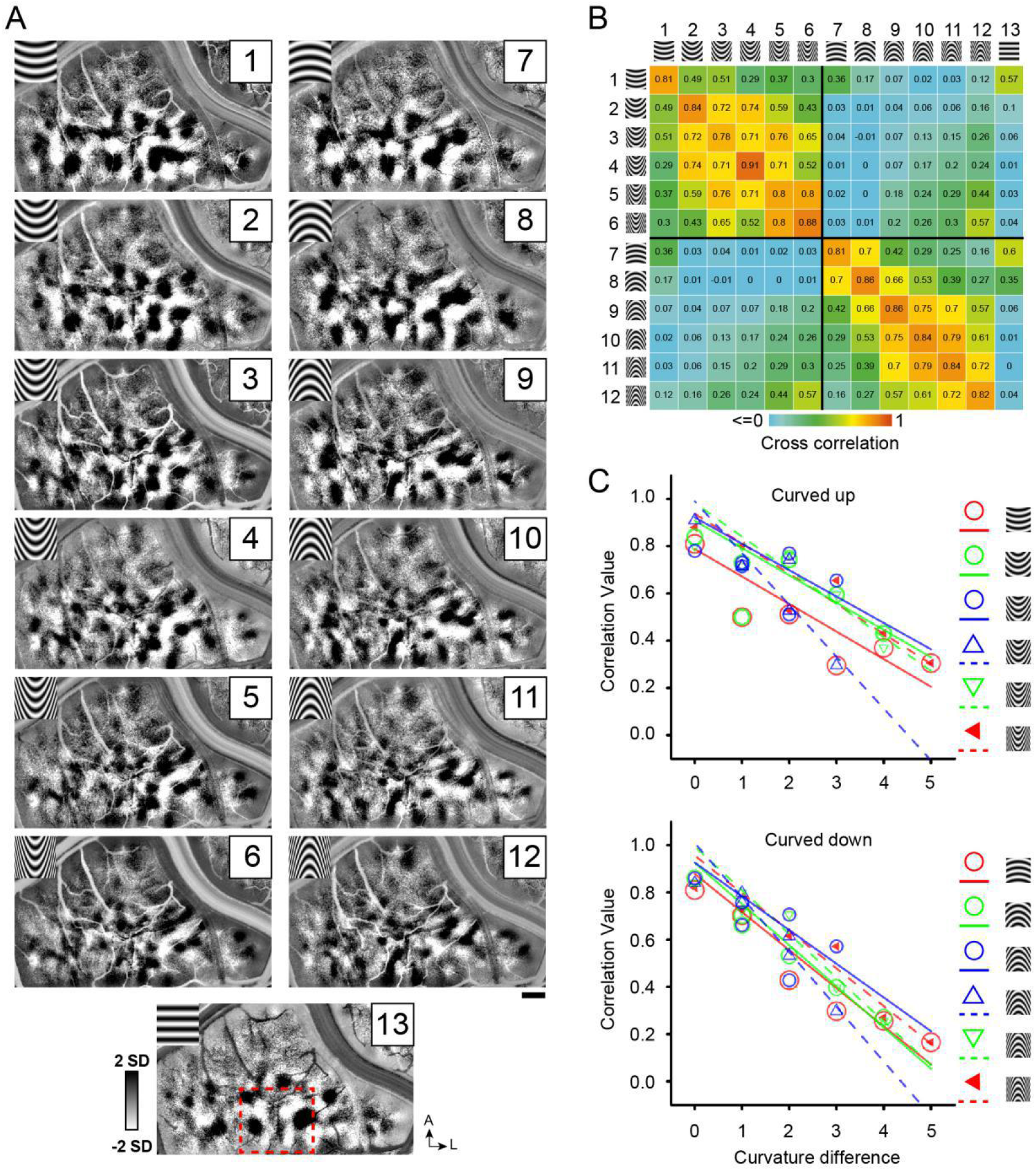
Systematic maps of curvature degree. A. Maps of different curvature degrees. Each map is one curvature minus sum of all straight gratings (Case 1). 1-6 and 7-12: upwards and downwards curvatures, respectively, from low to high curvature preference, 13: straight grating. Red dotted square marks the region that is further analyzed in Fig. 6A. B. Correlation values for pairs of curvature response maps (from A). Color bar: high (red) to low (blue) correlation values. Values between pairs of identical curvature degree (e.g. 1-1, 2-2) are calculated by correlating two maps, each obtained from half of the imaging trials. C. The more similar the curvature the greater the correlation value. X axis: curvature degree difference. Y axis: correlation value. Color symbols: correlation value for each curvature above with respect to its curvature degree distances (each fit with matching color line; see Table S3 for regression values). Scale bar, 1mm.

These distinctions are further reinforced by the finding that (Fig. 3A, Case 1 and Case 2) (1) curvature domains (white pixels: curvature vs. straight) contain subregions of high curvature preference (red pixels: high curvature vs. low curvature), (2) preference of these high curvature preference subregions (Fig. 3B, Case 1) for high (dark red lines) over low (light red lines) curvature, and negligible response to straight gratings (gray lines), and (3) the response magnitudes of these subregions gradually diminishes with curvature degree difference (Fig. 3C, dark to light lines plot responses of high to low curvature; top: upwards curvature, bottom: downwards curvature). In sum, these data suggest there are organized maps for curvature degree, that high curvature subdomains map within curvature domains (Fig.3A), and that curvature subdomains exhibit graded response to curvature degree (Fig.3B, C).

**Fig. 3.**
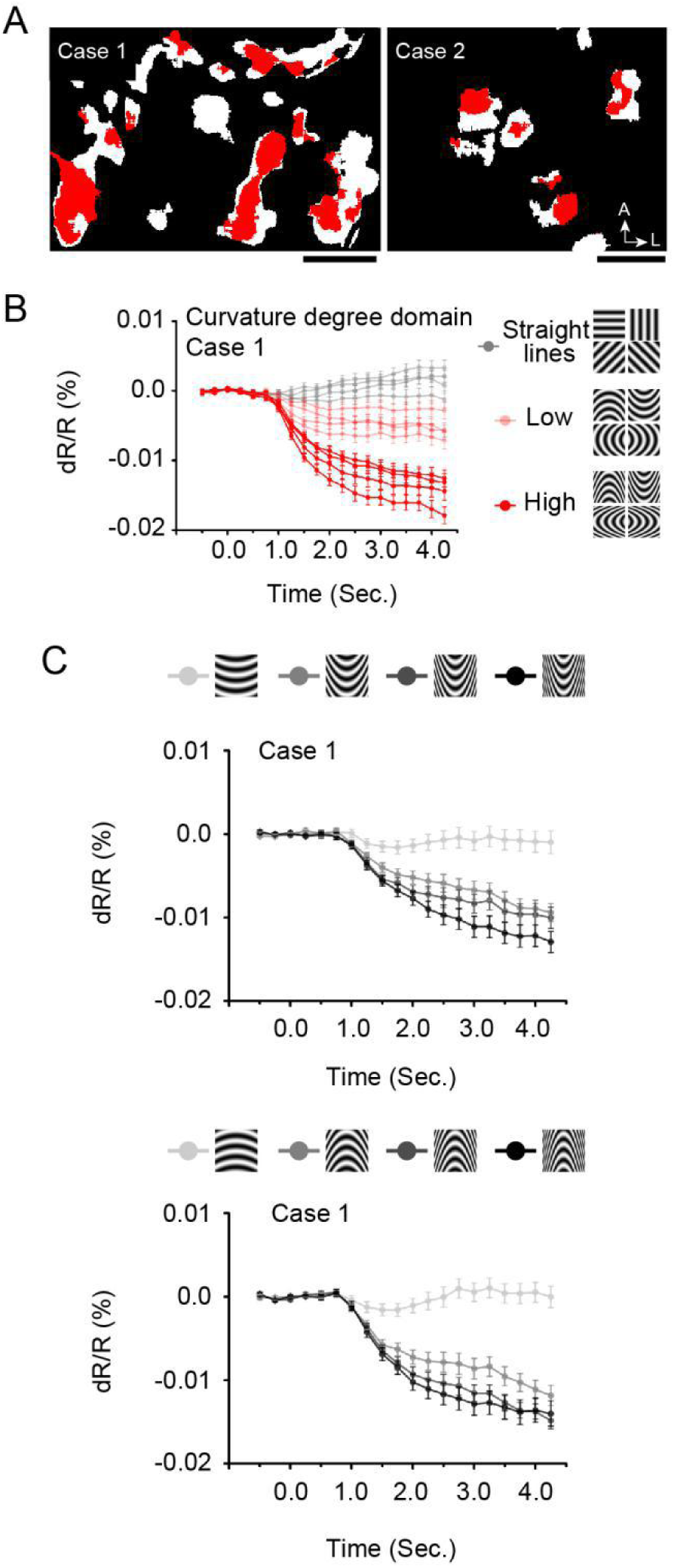
Subregions of high curvature preference. A. Curvature preferring pixels. Red pixels: high curvature preferring pixels (high curvature vs. Low curvature, a/b ratio =5 vs. 2, all 4 orientations, two-tailed t-test, p<0.01). White pixels: (curvature vs. straight). (Case 1: left, Case 2: right, two-tailed t-test, p<0.01). B. High curvature subdomains (red pixels in A, Case 1) prefer high curvature (dark red lines, 4 orientations, a/b ratio=5) over low curvature (light red lines, 4 orientations, a/b ratio=2), and straight lines (Gray lines, 4 orientations). C. Timecourses of response to 4 degrees of curvature (from high to low, darkest to lightest gray). Scale bar, 1mm.

#### Curvature orientation maps

In another experiment (Fig. 4, Case 2; Fig. S11C-D, Case 1), we used 4 *orientations of curvature* (curved upwards, downwards, leftwards, and rightwards), each at low curvature (1-4) and high curvature (5-8). Similar to maps for straight orientation (Fig. S10), we expected stronger correlation between maps of similar curvature orientation and lower correlation for dissimilar orientations. This expectation was supported by computed correlation indices for pairs of images (Fig. 4C). These correlation values illustrate that high correlation values (orange, yellow) occur only between curvature gratings of the same orientation (e.g. 1-5, 2-6), and low correlation values occur between curvatures of different orientations (green boxes in columns 1-8, e.g. 1-2, 3-4). The finding that the lowest correlation values occur between curved and straight gratings (columns 9-12) supports the distinctness of straight vs. curved contour representation. Similar results were obtained in a second case (Case 1, Fig. S11D).

**Fig. 4.**
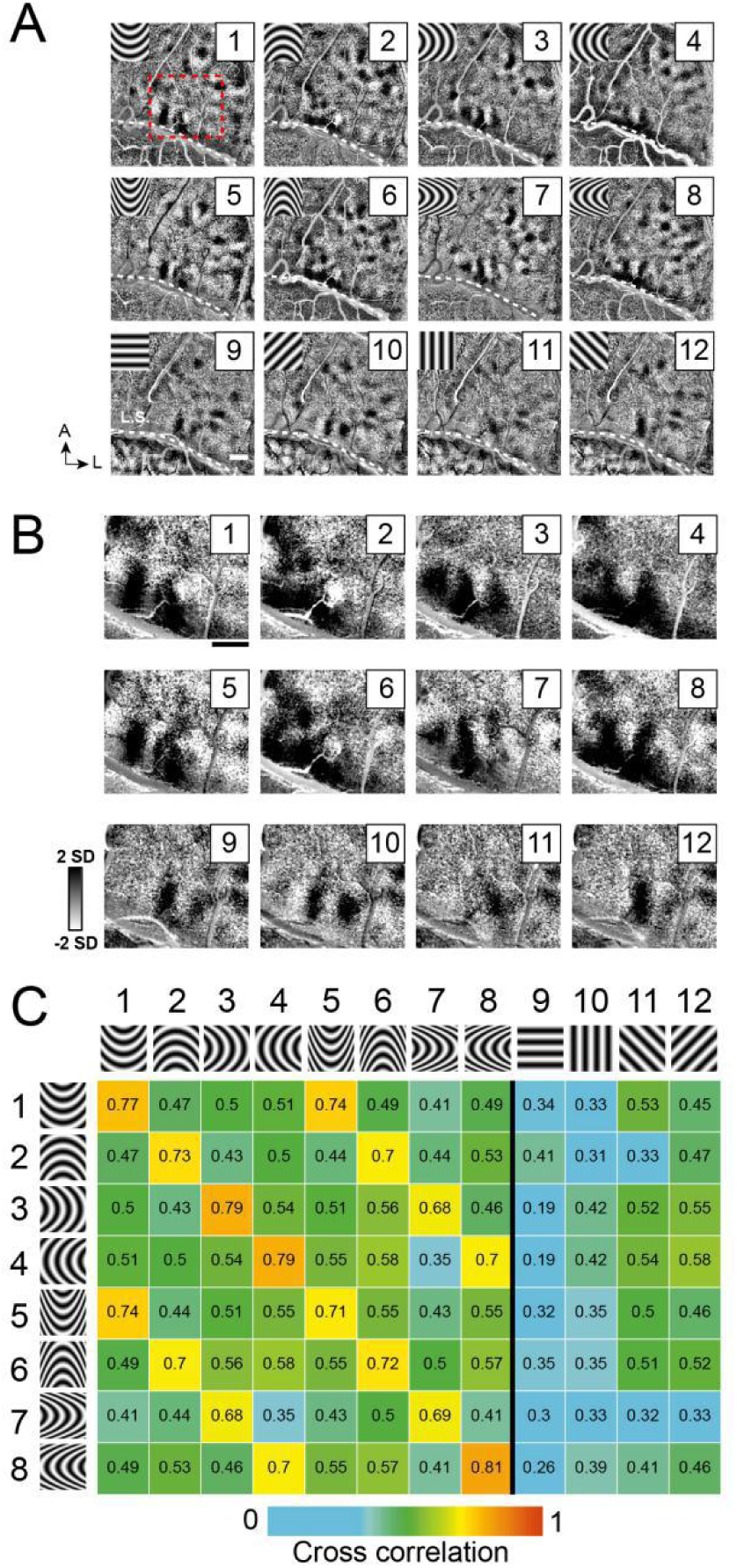
Systematic maps of curvature orientation. A. Imaging results of the responses to different curvature orientations. Maps of curvature minus sum of straight gratings (Case 2). 1-4: low curvature degree, a/b ratio=2, 5-8: high curvature degree, a/b ratio=5. 1,5: upwards. 2,6: downwards. 3,7: leftwards. 4,8: rightwards. 9-12: straight. B. Enlarged view of the cortical region outlined by red dotted box in A. C. Correlation values for pairs of maps (from A). Maps of similar curvature orientation have high correlation values, while those with different curvature orientation have low correlation values. Colors code correlation values (high, red, to low, blue, see color bar). Values between pairs of identical curvature orientation (e.g. 1-1, 2-2) are calculated by correlating two maps, each obtained from half of the imaging trials. Scale bar, 1mm.

To examine whether these curvature orientation maps exhibit any cue invariance, we conducted another experiment. In another case (Fig. 5, Case 3), we mapped 4 orientations of grating curvature (Fig. 5A) and then isolated the pixels preferring respective curvature orientations (red pixels). We then plotted the response timecourses of these curvature orientation selective pixels to curved gratings (Fig. 5B) and to *single curved lines* (Fig. 5C) presented at four orientations. We found that the preference of these pixels for single curved line orientation matched that for the curved grating orientation. For example, the pixels preferring the upwards curvature grating exhibited strongest response for upwards curved line (black lines in graphs in Fig. 5B and C). This curvature orientation cue invariance was also evident in a second case with two curvature orientations (upwards and downwards, Case 2, Fig. S12). These results reinforce the presence of subdomains selective for curvature orientation and provides additional support for curvature orientation cue invariance.

**Fig. 5.**
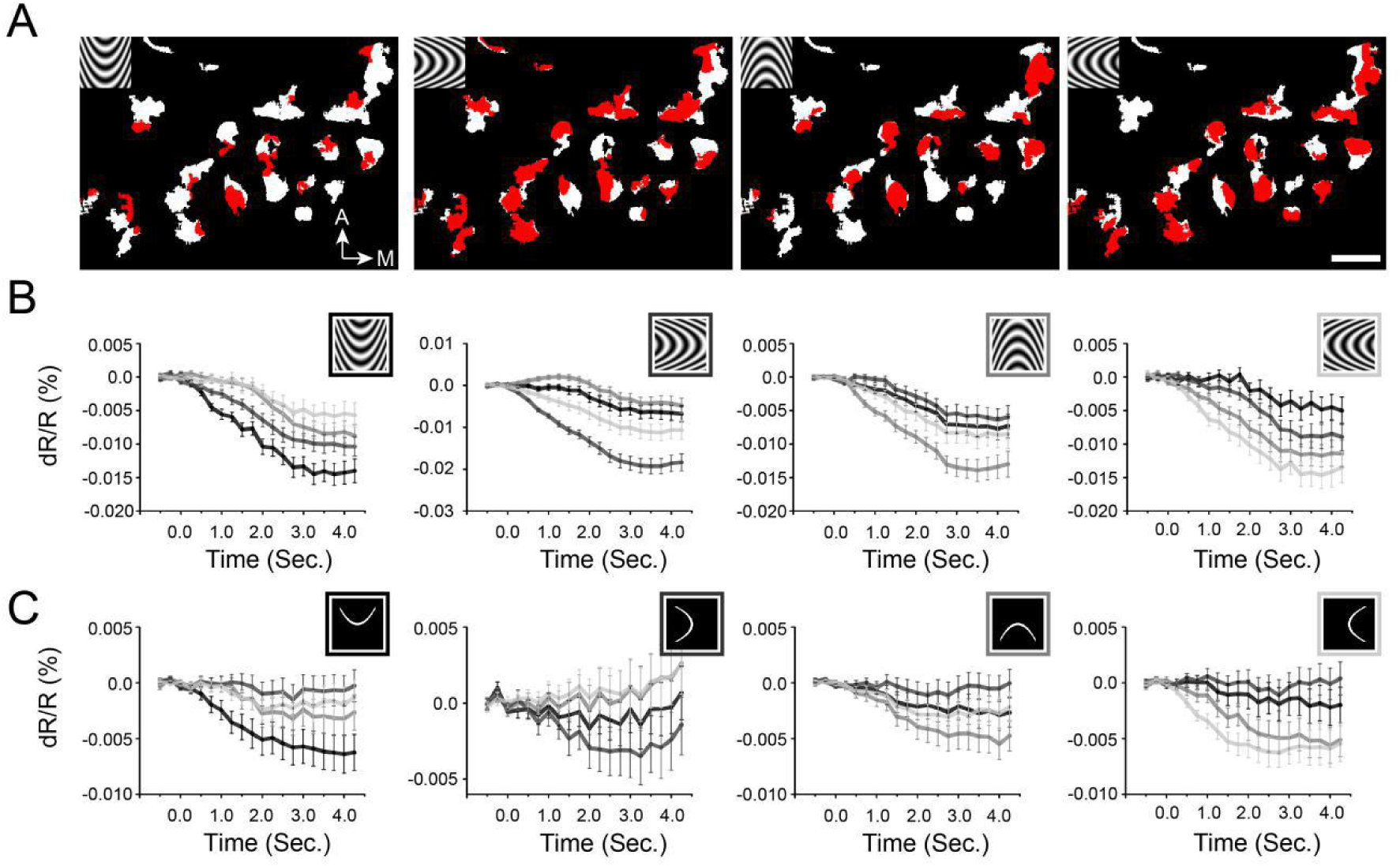
Curvature orientation domains are cue invariant. A. Curvature orientation maps. White pixels: curvature vs. straight (two-tailed t test, p<0.01). Red pixels: selective for different curvature orientations. B. Response timecourses of red pixels in maps above to curved gratings. C. Response timecourses of red pixels in maps above to single flashed curved lines. Black lines: response timecourses to upwards curvature. Dark gray lines: response timecourses to leftwards curvature. Gray lines: response timecourses to downwards curvature. Light gray lines: response timecourses to rightwards curvature. Comparison of B and C graphs show that the orientation of the best curved grating response matches that of the best curved line response (gray timecourses). Scale bar, 1mm.

### Organization of curvature representation

Given that there are indications of shifting maps for curvature degree and distinct maps for curvature orientation, we examined whether we could discern any systematic relationship between domains for straight orientation, curvature orientation, and curvature degree. We hypothesized that there is a gradual shift from straight to curved representation. As shown in Fig. 6A (Case 1), domains (left panels, Up: top row, Down: bottom row) are outlined by their preference from straight (blue) to high curvature (orange) (color code at top, images shown in Fig. S13A). In Fig. 6A (right panels), we demarcate the center of each domain with a color-coded dot (geometrical center). We observed a spatial progression from the center of the straight orientation domain (blue dot in center of blue dotted line) to low curvature domain (blue-oranges) to high curvature domain (orange); these colored dots are roughly connected by a line (shaded from blue to orange) to illustrate the progression (see Fig. S13). Four examples of such progressions for Up are shown in the top panel and four progressions for Down in the bottom panel. Note that progressions arising from a single orientation domain can lead to two different curvature domains (e.g. Up: leftmost orientation domain, Down: rightmost orientation domain). These results suggest some linkage between the representation of straight contours and curved contours.

**Fig. 6.**
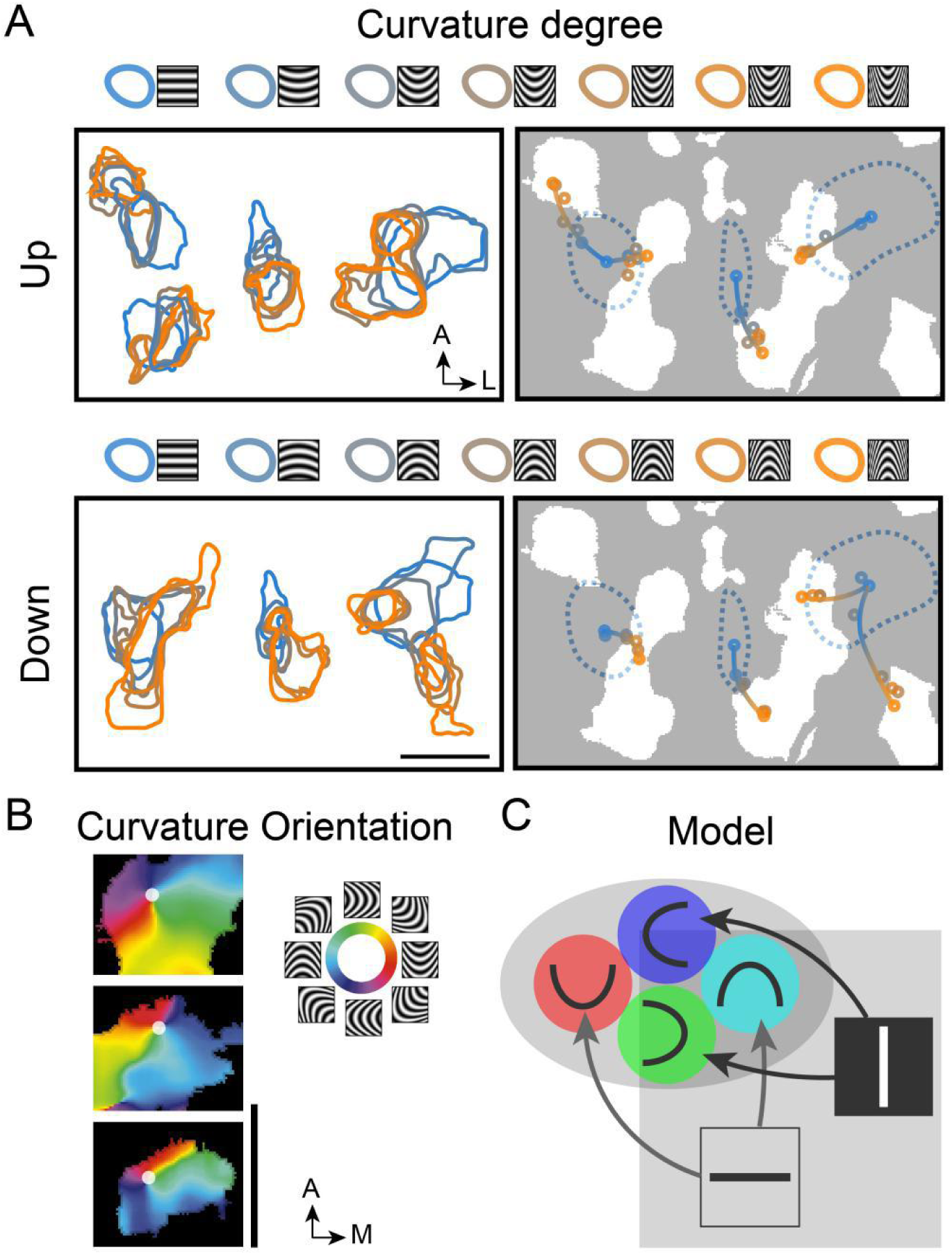
Functional organization of curvature in V4. A. Maps of progressions from straight to curved representation. Top row: responses to upwards curvatures (Up). Bottom row: responses to downwards curvatures (Down). *Left panels:* activated regions corresponding (two-tailed t test, p<0.01, curved vs. sum of straight) to respective curvature degrees are outlined by different colors. Color code (at top): high (orange) to low (blue). *Right panels:* Location of the activation center of each domain (indicated by colored dot). White regions: curvature domains. Blue dotted lines: horizontal orientation domains. Shifting progressions from straight orientation (blue dot) to low curvature (blue-orange dot) to high curvature (orange dot) are observed, as indicated by colored line (shaded from blue to orange). [Note that the two leftmost domain progressions in Up panel are associated with the same orientation domain but are separated for clarity.] More detail available in Fig. S13. Scale bar, 1mm. B. Maps of curvature pinwheel-like structures. Color-coded curvature orientation preference map (from Case 3). Colors represent curvature grating orientation within curvature domains, as shown by key on the right. White dot: pinwheel center. Scale bar, 0.5mm. C. Proposed organization of straight and curved contour representation in V4 orientation bands. Gray oval: curvature domains, Gray square: orientation domains. Colored circles: different curvature orientations. Squares in orientation domains: horizontal (black bar) and vertical (white bar) orientation domains. Arrows indicate progression from straight to low curvature to high curvature.

Another aspect of feature-specific maps is the presence of pinwheel-like progressions. Pinwheels are hypothesized to be landmarks of continuous ‘polymaps’ (for optimal representation of multiple feature spaces within a single area) and have been observed in both experimental data and computational models (21-23). Such structures have been previously observed in orientation maps and color maps in V1 (22, 24-26), V2 (27), and V4 (15, 17, 28) and more recently has been proposed as a fundamental unit of the orientation hypercolumn in V1 (26). To better evaluate this possibility, we conducted another experiment using a finer gradation of orientations (8 different orientations). As shown in Fig. 6B (from Case 3), we provide a few examples of curvature orientation organization reminiscent of previously reported pinwheel-like organizations. Finally, to further illustrate the distinction between curvature and straight orientation domains, we show curvature pinwheels and straight orientation pinwheels occupy distinct locations in the V4 map (Fig. S14).

While the overall spatial relationship between straight and curved features will take additional study to fully establish, given the data at hand, we propose the model shown in Fig. 6C. Our data suggest that straight (large gray square) and curved (large gray oval) regions in V4 co-exist and occupy roughly complementary territory. Within the curvature domains curvature pinwheels serve to represent different orientations of curvature, while curvature degree is mapped in a different axis, extending from the straight orientation domain to the curvature domain.

## Discussion

### Summary

In summary, we provide evidence for the existence of curvature domains in V4 and for a systematic map of curvature representation in V4. Maps for curvature are distinct from those for straight orientation, color, and high spatial frequency, and have the spatial appearance of organized functional maps. Our evidence suggests that curvature domains exhibit selectivity for degree of curvature (low to high), one which maps in a spatially shifting manner in V4. We also find domains selective for orientation of curvature. The presence of pinwheel-like organizations characteristic of other functionally organized maps in visual cortex are also consistent with a systematic organization. Furthermore, the invariance of response across both curved gratings and single curved lines indicate that these are high order responses consistent with curvature encoding.

### Novel finding

Milestones have been achieved in the past decades in understanding of curvature coding in the cortex, (1) Electrophysiology. Electrophysiological recordings in V4 have found evidence for neurons preferring curved over straight contours (9, 11, 40, 41). There is also substantial evidence supporting a neural population code from which object shape can be reconstructed or decoded (30). Marge Livingstone’s lab reported using multi electrode arrays that there is a strong correlation between curvature selectivity and end-stopping in V1, V4, and inferotemporal cortex, 20). (2) Neuroimaging: As demonstrated by fMRI, Leslie Ungerleider’s group reports there are 3 curvature preference areas (roughly several mm’s to > 1cm in size) in macaque V4/TEO/TE which form a curvature processing network (42). (3) Modelling. There are multiple modelling studies examining how V4-like curvature units contribute to shape representation (12, 13). However, these pioneering studies do not have the resolution to demonstrate column-based (submillimeter) functional organization in these areas. No previous optical imaging study has reported functional organization for curvature in V4. Thus, to our knowledge, there is no study that has reported the functional organization for curvature in V4. Furthermore, no study has examined the relationship of curvature domains with other types of functional domains. Finally, no study has provided evidence suggesting that curvature is a basis for a hypercolumn in V4.

### Unlikely to be high spatial frequency maps

We consider the possibility that the curvature responses are actually responses to the high spatial frequency aspects of the curvature gratings. Based on curvature stimuli used in Ponce et al 2017 (20), we designed curvature gratings composed of ellipsoid curves repeated sinusoidally. As shown in Fig. S6, different stimuli with different degrees of curvature created in this fashion have common spatial frequency along the center axis. Towards the flanks of the stimulus, especially for higher curvature, the grating has higher SF content than that on the central axis. However, several observations make our results inconsistent with the hypothesis that these results are due primarily to these high SF aspects of the stimuli. (1) High spatial frequency maps in V4 (first reported by Lu et al., 2018, 19) differed from curvature maps. On average, less than 15% overall from three cases exhibited overlap between high spatial frequency domains and curvature domains. (2) Similar maps were obtained to gratings with different degrees of curvature (Fig. S7). If the results were due primarily to spatial frequency content, the maps resulting from high and low curvature gratings should differ. (3) Similar maps were obtained to gratings and single lines of the same curvature (Fig. 1K, L, Fig. 5), although gratings and single lines have different spatial frequency contents. (4) Scrambled maps, scrambled stimuli which are composed of small patches of the curvature stimuli but randomly arranged in space, failed to produce curvature maps. Thus, while we have not excluded all possibilities, we believe that the bulk of the evidence provides a consistent view, one which has never previously been demonstrated. As with many new findings, there will be more studies to follow.

### Population coding

Previous single unit studies of curvature response in V4 have found that single neurons are tuned to particular curvatures and particular orientations of curvature (9-11). However, single neurons can respond to different curvatures with the same firing rate (e.g. the same degree of curvature at slightly different location or orientation) (11). Consequently, as previously concluded, encoding of shape in V4 is unlikely to be accomplished by single neurons. Instead, many studies support a population based representation of shape (29-34). Each neuron involved in this coding process contributes to different components of the shape. Based on the weighted integration of participating neurons, the location and curvature information of a shape may be accurately described (30). This ‘population code’ has been highly influential in our conception of how shape is encoded in the brain. However, these population results were acquired through neuronal recordings acquired without knowledge of functional domain location.

In contrast to previous studies which sampled single neurons at a time, we aimed to understand the population response of all the points (via simultaneous optical imaging of thousands of pixels) within a large visuotopically appropriate location in V4. Using optically imaged topographic mapping methods, we carefully identified the cortical territory responsive to our stimulus (Fig. S2) and matched the sizes of our stimuli (4°) to typical receptive field sizes in the imaged region (9, 35). *Our findings are consistent with previous population based concepts, but introduce a systematic spatial view of curvature representation. Our study has revealed (1) that multiple domains are involved in the representation of single curvature gratings and single curved lines (Figs. 2 and 4), (2) that specific sets of domains represent different aspects (degree or orientation) of curvature, and (3) that the category (degree, orientation) of curvature can be classified from this representation (Figs. 1L, 4 and 5)*. These novel findings have not been revealed by electrophysiological recordings and suggest that, on average, the neuronal population code is spatially organized within a columnar population code.

### Cortical modules represent key transformational stages in the visual hierarchy

As we hypothesized a systematic continuum of curvature representation within V4, we designed stimuli that spanned curvature space with respect to curvature degree and curvature orientation. Instead of using shapes with complex structures such as hyperbolic/polar gratings (6) or forms composed of multiple corners and lines (30, 36), we used relatively simple curvatures. The gradations of curvature degree across our stimulus set revealed that response preference is spatially and continuously mapped in V4 (from straight to highly curved, Fig. 6A, and across different orientations, Fig. 6B). As evident from other cortical functional maps (15-17, 22, 24-26), this continuum of parameter representation (a la orientation maps in V1) suggests a common transformation of information representation in the cortex. In this case, the common transformation is curvature representation and is embodied in a module we term the curvature domain.

Why does it matter that the representation is modular and that it is systematically mapped? We suggest that it is modular organization that identifies the key functional transformation central to a particular cortical stage and which, in turn, reveals the functional identity of nodes within the computational visual cortical hierarchy. Here, we provide a brief explanation. Similar types of responses can be found in many areas of the brain. For example, each of V1, V2, and V4 contain orientation selective domains; however, orientation selective domains are categorically different in V1, V2, and V4. Whereas those in V1 are responsive to contrast-defined orientation, those in V2 have cue-invariant orientation selectivity and represent a higher order abstraction of orientation (5, 37, 38). We suggest that orientation selective domains in V4 are at yet a higher level. In fact, we hypothesize that they are not ‘orientation selective’ in either the V1 sense or the V2 sense, but actually represent an integration of lower order orientation inputs to generate representation of straight curves, one which lies within a continuum of low to high curvature representation.

We thus suggest that curvature domains in V4 represent the ‘third’ computational stage of processing in the shape processing hierarchy. This stage is replete with curvature degree progressions and pinwheel-like curvature orientation organization, providing a representational architecture for a ‘curvature hypercolumn’ in V4. Stated more broadly, whereas many types of single neurons may be found in any cortical area, modules and hypercolumns reveal the true signatures of a cortical area’s computational role.

## Materials and Methods

Data was acquired from 3 hemispheres of 2 adult macaque monkeys (one male and one female, Macaca mulatta). All procedures were performed in accordance with the National Institutes of Health Guidelines and were approved by the Zhejiang University Institutional Animal Care and Use Committee.

### Animal preparation

Chronic optical chambers were implanted above the area V4d containing lunate sulcus, superior temporal sulcus (see Fig. 1A, B) as described previously (16). The only difference is that we used transparent glass instead of a nylon chamber. The eccentricity of the visual field corresponding to the exposed V4/V2/V1 was 0-5°. Following the craniotomy surgery, optical images were collected during which basic functional maps as well as curvature responses of V4 were obtained. Monkeys were artificially ventilated and anesthetized with propofol (induction 5-10 mg/kg, maintenance 5-10 mg/kg/hr, i.v.) and isoflurane (0.5-1.5%). Anesthetic depth was assessed continuously via monitoring heart rate, end-tidal CO_2_, blood oximetry, and eeg. Rectal temperature was maintained at around 38C°. Animals were paralyzed (vecuronium bromide, induction 0.25 mg/kg, maintenance 0.05-0.1 mg/kg/hr, i.v.) and respirated. Pupils were dilated (atropine sulfate 1%) and eyes were fitted with contact lenses of appropriate curvature to focus on a stimulus screen 57 cm from the eyes.

### Visual stimuli for optical imaging

Visual stimuli were created using ViSaGe (Cambridge Research Systems Ltd.) and displayed on a calibrated 27-inch monitor (Philips 272G5D) running at 60 Hz refresh rate. The luminance for white stimuli was 206.52 cd/m^2^ and black was 0.50 cd/m^2^. Full-Screen visual grating stimuli were used to locate color preference domains in V4. Red/green isoluminance and black-white sine-wave drifting grating stimuli were presented at two different orientations (45° and 135°) with the same spatial frequency (0.5 or 1 cycles/°), temporal frequency (2 or 4 Hz) and mean luminance level (see Fig. S3A). To acquire orientation maps and spatial frequency (SF) maps, gratings with 4 different orientations (0°, 45°, 90°, 135°, see Fig.S3A) and two different SFs (0.5, 4 cycles/deg) were presented.

Curvature stimuli were designed based on the formula of ellipse, X^2^/a^2^ + Y^2^/b^2^ = 1, where a and b represent the length of the long axis and short axis respectively. Different a/b ratios were used (2, 5, See Fig. 3B; 1, 4, 7, 10, see Fig. 3C) to create different curvature degrees. The ellipses (see Fig. S3A) were cropped and duplicated to generate a curvature grating template. SF (along the long axis) of the curvature stimuli was varied by changing the spacing of the curves. The curvature grating was drifted by moving the stimulus window along the curved grating template (see Fig. S3B). Drifting straight gratings were also created by this process and had the same luminance, drift speed, and spatial frequency (at center axis of curvature) as curvature stimuli (Fig. 1C). For curvature maps, the position of the stimuli on the monitor was first determined (Fig. S2). As a control for curvature, scrambled versions of a curvature grating were created by dividing the grating into 64 subunits and randomly rearranging the locations of these subunits.

#### Retinotopic mapping

For placement of stimuli on the monitor, we mapped the retinotopy of V4 using a series of 0.2° width horizontal or vertical dashed lines (SF=1 cycles/°) located at different positions to determine the stimulus center (locating the stimulus center at the imaged V4 center, *16*, see Fig. S2). As the receptive field sizes of V4 neurons at our recording position are around 3-5 degrees (*35*), our stimuli were set to 4 degrees and presented monocularly during the experiment. Flashed curved lines were also 4° in size (for each line, the length was 4°). The center of straight lines or the vertex of curved lines were fixed at the position that we chose during the retinotopic mappings. The stimuli were presented at 4 or 8 Hz interleaved with black screen.

### Optical Imaging

The brain was imaged through the implanted glass. Images of cortical reflectance changes (intrinsic hemodynamic signals) corresponding to local cortical activity were acquired (Imager 3001, Optical Imaging Inc., German town, NY) with 632 nm illumination. Image size was 1080×1308 pixels representing 14.4×17.4 (case 2) or 8.7×10.5 (case1 and 3) mm field of view (see Fig. S1). Visual stimuli were presented in a random order. Each stimulus was presented for 3.5 (for color and high spatial frequency stimuli) or 4.5 (curvature and corresponding straight stimuli) seconds. The imaging data were stored in a block fashion. Each block contained the imaging data recorded from all the stimulus conditions (presented one time). For functional domains, each stimulus was presented at least 30 times. For retinotopic mapping (Fig. S2), each stimulus was presented 10 times. Imaging started 0.5 s before the stimulus onset and ended till the stimulus offset with a sampling rate of 4Hz.

### Data analysis

#### Functional maps

With the following formula, 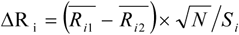, we assessed the response differences between two comparison groups. 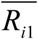 and 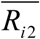 are the mean dR/R values (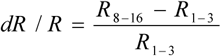, R_8-16_ is the averaged response from frame 8 to 16, R_1-3_ is the averaged response from frame 1 to 3) in the two compared conditions of pixel i respectively, N is the number of trials, and S_i_ is the standard deviation of (*R*_*i*1_ − *R*_*i*2_). Color preference maps were obtained by comparing red/green and white/black grating images, spatial frequency maps by comparing high and low spatial frequency images, and orientation maps by comparing two orthogonal orientation images (0° vs. 90° or 45° vs. 135°). For curvature maps, we compared the sum of curved gratings with the sum of straight gratings (four orientations: 0°, 45°, 90°, 135°). For single condition maps, we compared the selected condition with the averaged of straight gratings (four orientations: 0°, 45°, 90°, 135°). Maps were high-pass filtered (Gaussian filter, 10 pixel diameter) and low-frequency noise was reduced by convolving the map with a 100-150 pixel diameter circular filter and subtracted from the original maps. The orientation preference angle maps were calculated based on the single condition maps, and each pixels were assigned with a unique color to represent the preferred orientation (*39*).

#### Locate the positions of functional domains

Functional domains were identified by selecting the pixels with significant dR/R difference (two-tailed t test, p<0.01) between two comparison conditions (color domains, red/green versus white/black, ΔdR/R<0; high spatial frequency domains, high spatial frequency versus low spatial frequency, ΔdR/R<0; 0° orientation domain, 0° versus 90°, ΔdR/R<0; 45° orientation domain, 45° versus 135°, ΔdR/R<0; 90° orientation domain, 90° versus 0°, ΔdR/R<0; 135° orientation domain, 135° versus 45°, ΔdR/R<0; Curvature domains, curved gratings versus straight gratings, ΔdR/R<0). In addition to t-test, pixels that belong to curvature domains were further assessed by one-way ANOVA (curved stimuli versus straight stimuli, p<0.05 with post–hoc Tukey Kramer correction) test. Domain size was calculated as size = πR^2^, where area size is the size of each separated patch (pixel number× area of each pixel), R is the average of long and short axis of patch. Patches smaller than 0.2mm (diameter) were excluded from this analysis, as the reliability of these small patches is less secure (see Fig. S8). For each activated region, the geometrical center was calculated as its activation center (see Fig. S6C).

#### Timecourse

For time courses, the gray value of each pixel was calculated first using the following functions: dR/R=(Fx-F0)/F0, where F0 is the average reflectance value of the first two frames (taken before visual stimuli onset), Fx is the reflectance value corresponding to frame X (X= 1, 2, 3, …). Pixels with significant response changes (p < 0.01, two-tailed t test) between the two comparison conditions were included. The response time courses were obtained by averaging the values of these selected pixels (see Fig. 1, 3, 5). In order to calculate the response amplitudes of different functional domains in different stimuli, the last 10 points in corresponding timecourse from different cases were averaged. These data were also used to test whether there is significant response difference between two conditions (see Fig. 1L).

#### Similarity

To evaluate the similarity between two single condition maps, we extract the responses in the imaged area V4 (regions that could be activated by curvature stimuli during the experiment) and calculated the correlation coefficient values between the two response maps (see Fig. 2 and 4). To evaluate the relationship between curvature degree difference and response similarity, we use the Regression function from Matlab. In this analysis, we use the condition number to represent the relative curvature degree of each curvature (see Fig. 2). The curvature degree difference is based on the relative distance between two curvature degrees.

#### Statistical Analysis

For all statistical comparisons of functional maps, we use two-tailed t test or ANOVA. We used the Wilcoxon rank sum for the comparisons between response amplitudes in two conditions when data failed to show a normal distribution.

## Acknowledgments

We thank Robert M. Friedman for helpful comments.

## Author contributions

J. M. H. carried out most of the experiments and analyzed the data. X. M. S. and Q. N. W. helped with experiments and data analysis. A. W. R. and J. M. H. conceived the project and wrote the manuscript.

