## Supplemental data for "Curvature domains in V4 of Macaque Monkey"

### Supplementary Materials

**Fig. S1.**

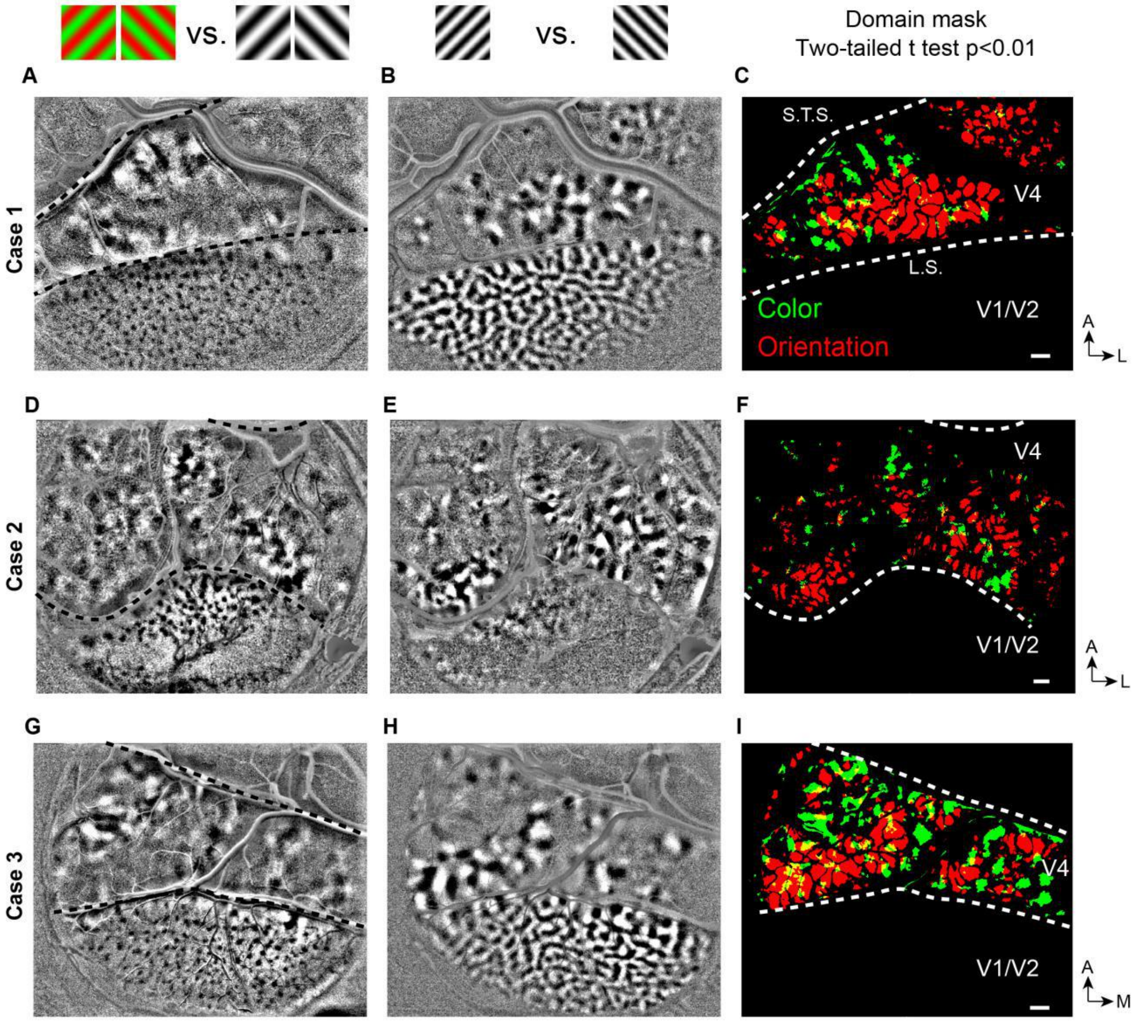


**<u>Fig. S1.</u> Large field of view functional maps in Cases 1-3.** Color (left column) and orientation (middle column) maps in V1, V2, and V4. Third column: overlay of color preferring (green) and orientation selective pixels (red). White dotted lines: lunate sulcus (LS) and superior temporal sulcus (STS). For all maps, the locations of functional domains were determined by t-value maps (t-map, two-tailed t test, p<0.01) which were calculated by comparing, pixel by pixel, the responses between two different conditions (A: color vs. achromatic, B: 45° vs. 135°). Scale bar: 1 mm.

**Fig. S2.**

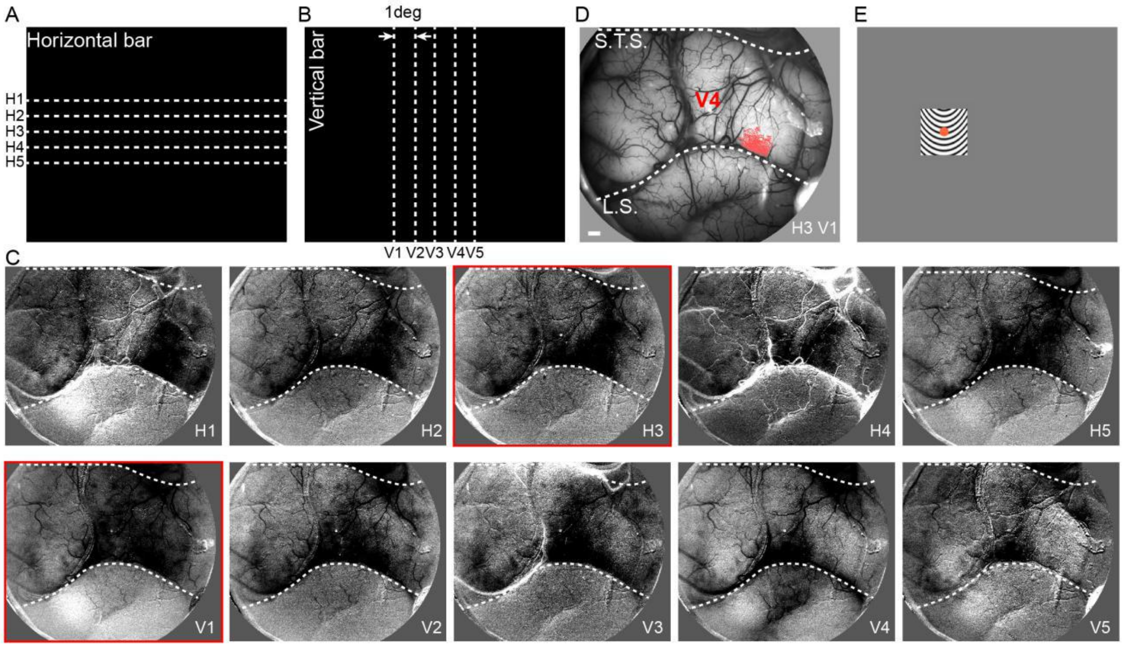


<u>**Fig. S2.**</u> **Retinotopic mapping in V4.** A, B. Visual stimuli used for retinotopic mapping. C. Cortical responses to the visual stimuli marked by different numbers in A and B. These maps were used to guide the placement of curvature stimuli (as shown by red dot in E). D, Red pixels: Cortical location corresponding to the center of the curvature stimulus (i.e. pixels that are significantly activated, two-tailed t test, $p<0.01$, for both the horizontal and vertical positions, in this example, H3, V1). E, Corresponding stimulus position on the screen. Scale bar: 1 mm.

**Fig. S3.**

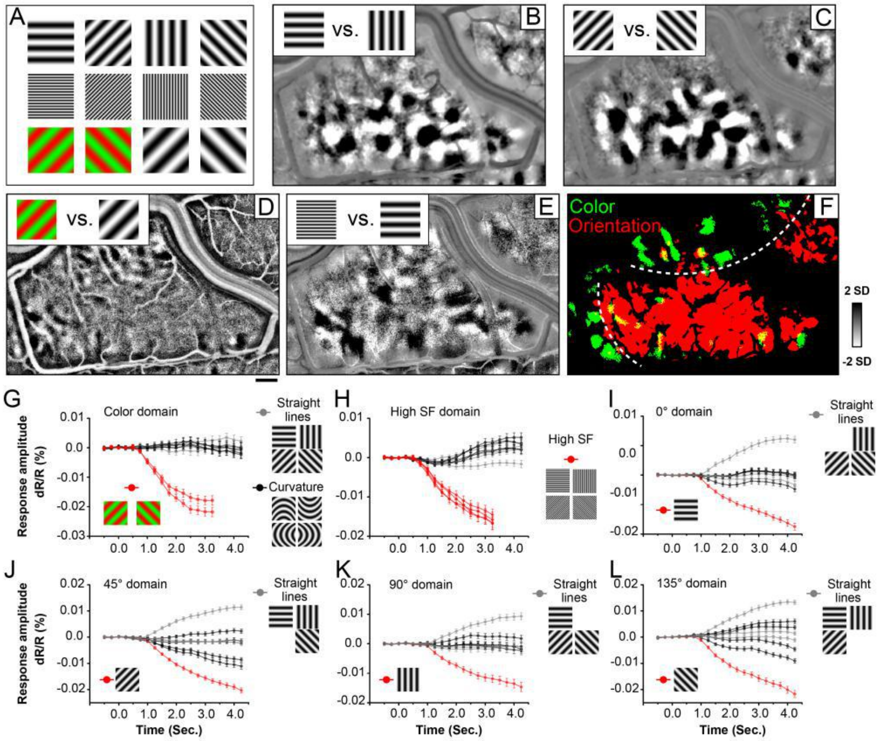


**<u>Fig. S3.</u> Functional maps in macaque monkey V4.** A. Visual stimuli. B-L: Case 1. B-C. Orientation maps. D. Color preference map. E. High spatial frequency preference map (4 cycle/deg vs. 0.5 cycle/deg). F. Overlay of color map (green pixels: red/green gratings versus black/white gratings, two-tailed t test, $p<0.01$) and orientation map (red pixels: orientation selective pixels from B and C, two-tailed t test, $p<0.01$). Yellow pixels: regions of overlap. White dashed lines in F: approximate borders between color and orientation bands. Scale bar: 1 mm. G-L. Response time courses of different functional domains. Red lines: preferred stimuli for each domain type. Gray lines, black lines: non-preferred stimuli (see insets next to each graph). G: color domains. As expected, color domains exhibited robust response to isoluminant color gratings (red line) but weak response to achromatic gratings (gray lines: straight gratings, black lines: curved gratings). H: high spatial frequency domains. Similarly, high spatial frequency preference domains responded strongly to achromatic gratings of high spatial frequency (red lines) but poorly to low spatial frequency gratings (either straight, gray lines, or curved, black lines). Note that, neither color domains nor high spatial frequency domains exhibited stronger responses to curvature stimuli. (color domains, averaged $dR/R=6.8\times10^{-6}$; high spatial frequency domains, averaged $dR/R=1.5\times10^{-6}$). I-L: orientation domains. As expected, straight orientation domains (I-L) exhibited strongest response to gratings of their respective optimal orientations (red line), relative suppression to gratings of orthogonal orientation (top gray line), and weak responses to other straight orientations (other gray lines). In comparison, they displayed relatively little response to curved gratings (black lines) (Fig. S3I, Wilcoxon test, $p<10^{-5}$ for each of the four optimal orientation responses compared to response to all 4 curvatures summed, see Fig. S3J-L for responses of 45º, 90º, 135º domains). These analyses indicate that curvature processing is not subserved by color or high spatial frequency domains. Preferences of orientation domains and curvature domains were distinct.

**Fig. S4.**

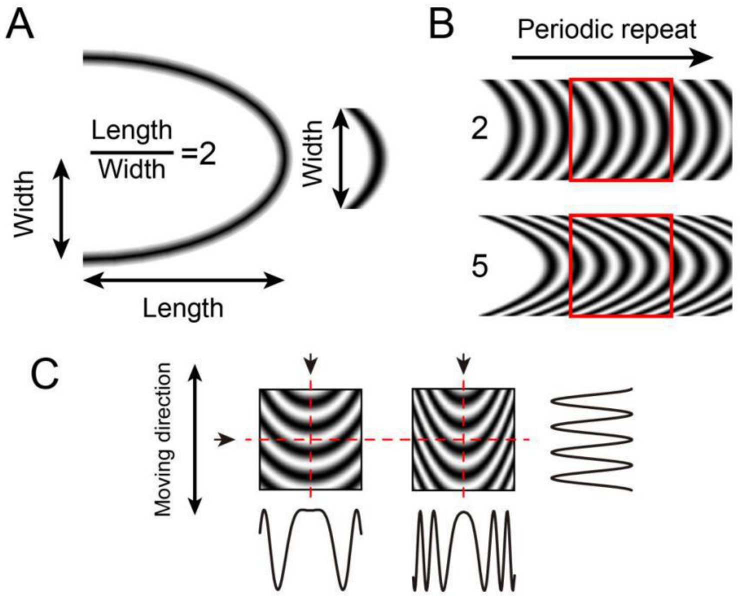


**<u>Fig. S4.</u> Curvature grating stimuli.** A. Calculation of curvature index. Width is half of the full width. B. Constructing curvature grating by periodic repeat. One low (index=2) and one high (index=5) curvature grating shown. C. Luminance profiles of different stimuli in different directions. The profiles at the vertical axis (arrows at top, profile at right) of the two stimuli are similar, but differ in the horizontal axis (arrow at left, profiles below). The higher the curvature degree, the higher the spatial frequency towards the edge of the stimulus.

**Fig. S5.**

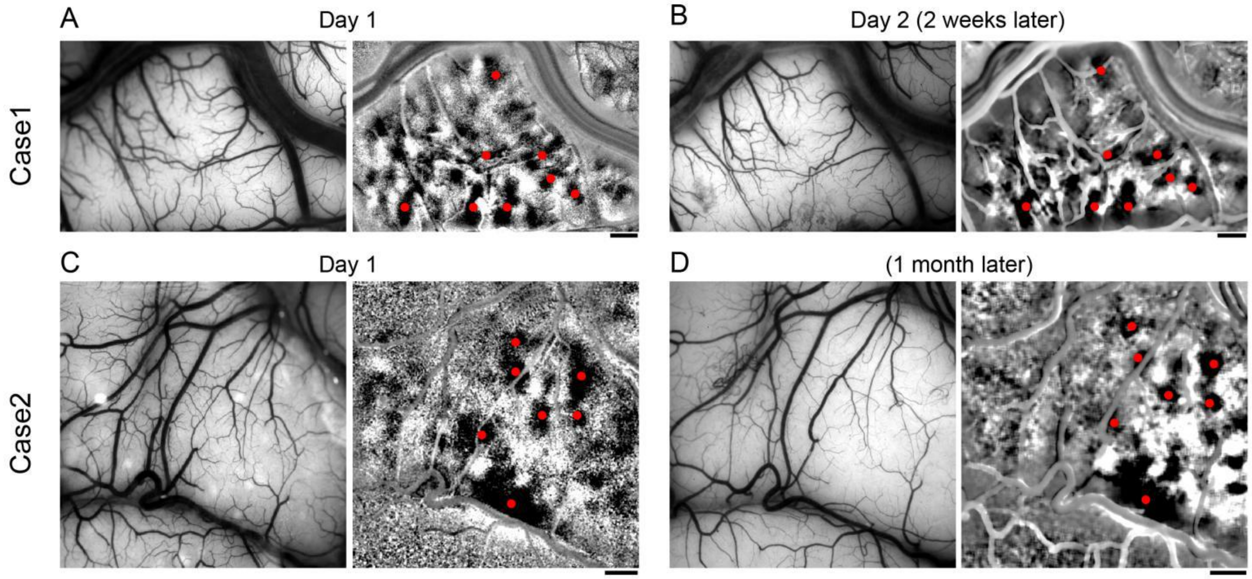


**<u>Fig. S5.</u> Imaging of curvature responses in the same cortical region (from two cases) in different days.** Maps of all curvature minus all straight. A and B are from the same imaging chamber, C and D are from the same chamber, imaged on different days. Red dots: corresponding curvature domains imaged on different days. Scale bar: 1 mm.

**Fig. S6.**

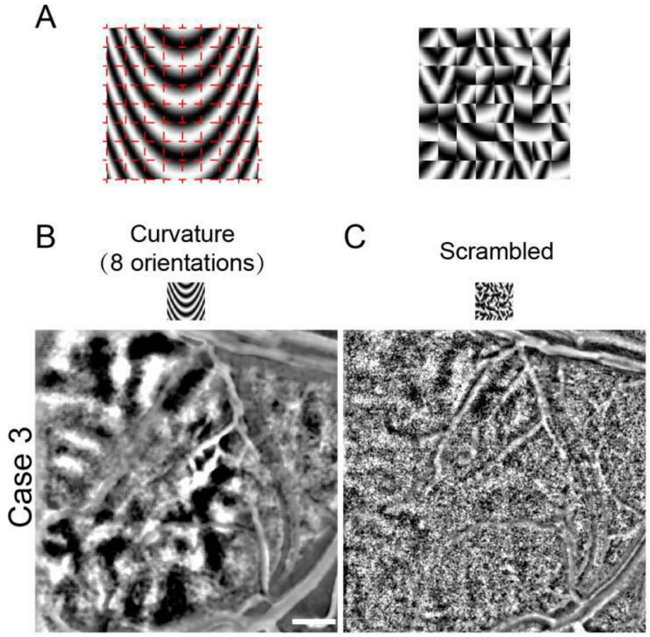


**<u>Fig. S6.</u> Imaging of curvature responses and scrambled map in the same cortical region (from Case 3).** A. Scrambled version of a curvature grating created by dividing the grating (each frame) into 64 subunits and rearranging the subunits randomly. B. Curvature map (curvature vs. straight). C. Scrambled map (Scrambled vs. straight). Scale bar: 1 mm.

**Fig. S7.**

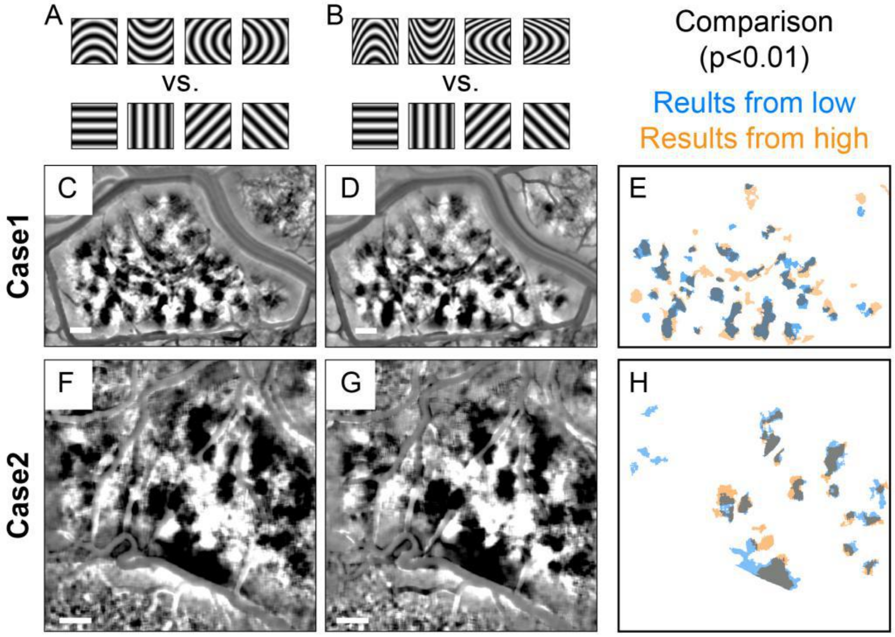


**<u>Fig. S7.</u> Curvature maps are not dependent on SF of curvature stimuli: curvature degree.** A-B: Stimuli with low (A, Length/width=2) vs. high (B, Length/width=5) curvature degrees.

These are compared with straight orientation gratings of the same SF. C-G: subtracted curvature maps for Case 1 (C,F) and Case 2 (D,G). E, H: Overlay of low vs. high T maps ($p<0.01$). Blue: low SF. Gold: high SF. Scale bar: 1 mm.

**Fig. S8.**

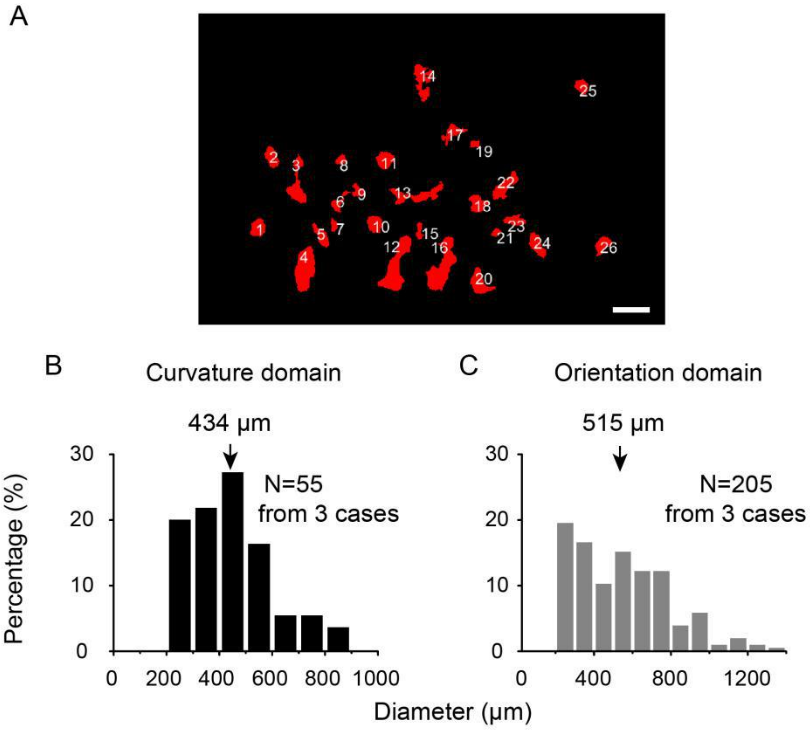


**<u>Fig. S8.</u> Curvature domain size.** A. Curvature domains in Case 1 (two-tailed t-test, $p<0.01$). Each domain is numbered. Scale bar, 1mm. B. Distribution of curvature domain diameters from three Cases 1-3 (n=55). C. Distribution of straight orientation domain diameters from Cases 1-3 (n=205). Domain size was calculated as size=$\pi R^2$, where area size is the size of each separated patch (pixel number× area of each pixel), R is the average of long and short axis of patch. Patches smaller than 0.2mm (diameter) were excluded from this analysis, as the reliability of these small patches is less secure.

**Fig. S9.**

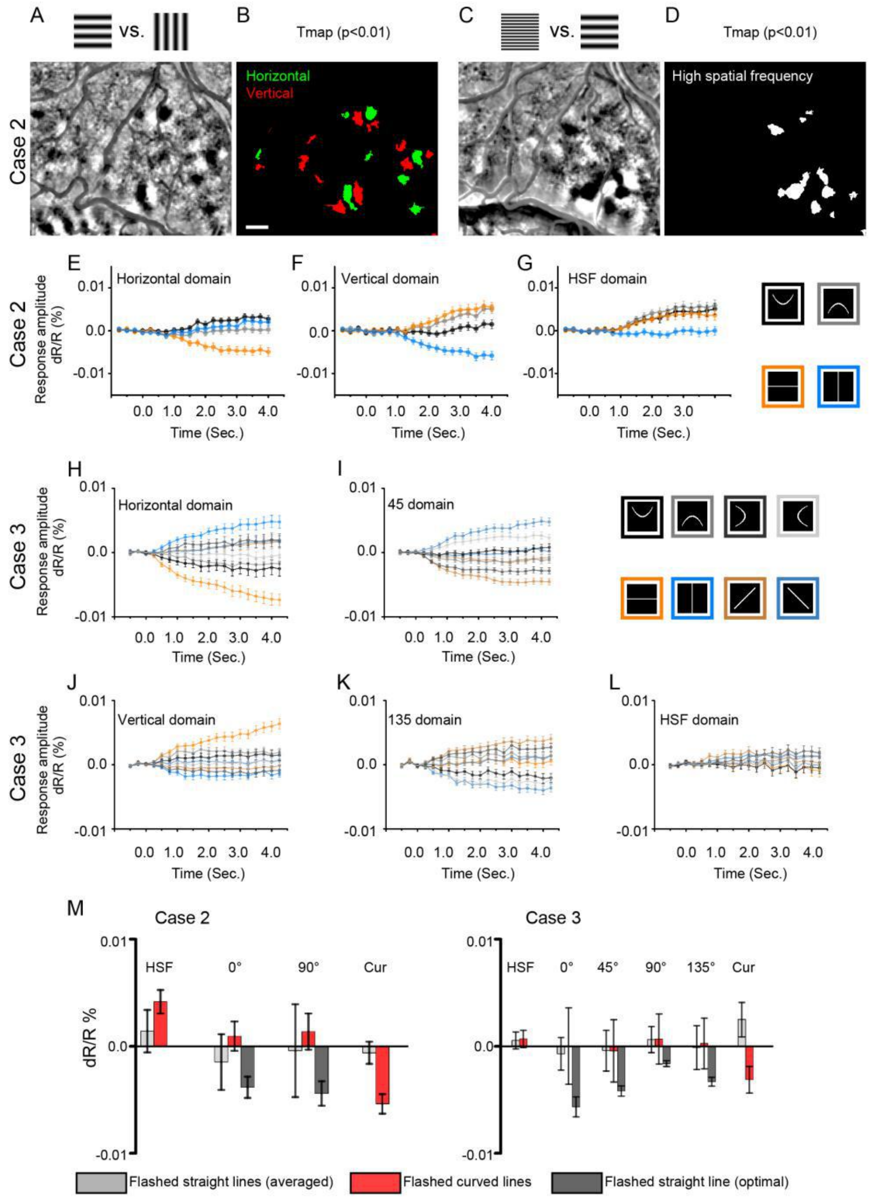


**<u>Fig. S9.</u> Comparisons of responses to single curved and straight lines in orientation and high spatial frequency domains.** A-G: Case 2. A. Orientation map. B. T-map of A (p<0.01). Green: horizontal; Red: vertical. C. Spatial frequency map [4 vs. 0.5 cycle/deg, with four different orientations, 0, 45, 90, 135]. D. T-map (p<0.01) of C. E-G. Response timecourses of horizontal (E), vertical (F), and high spatial frequency (G) domains in response to flashing of single curved lines (Black: upwards, Gray: downwards,) and single straight lines (Orange: 0, blue: 90). H-L: Case 3. Response timecourses of horizontal (H), 45(I), vertical (J), 135 (K) and high spatial frequency (L) domains in response to flashing of single curved lines (Black: upwards, Dark gray: leftwards, Gray: downwards, light gray: rightwards) and single straight lines (Orange: 0, Dark orange: 45, blue: 90, dark blue: 135). Compare with graphs in Figure 1K. Scale bar: 1 mm. M. Responses for high spatial frequency (HSF), orientation (Case 2: 0º, 90º; Case3: 0º, 45º, 90º, 135º), and curvature (Cur) domains in Case 2 (left) and Case 3 (right) to flashed lines. Light gray:

straight lines. Red: curved lines. Gray: Optimal straight lines. Error bars: SEM (timecourses in E-L), SD (histogram in M).

**Fig. S10.**

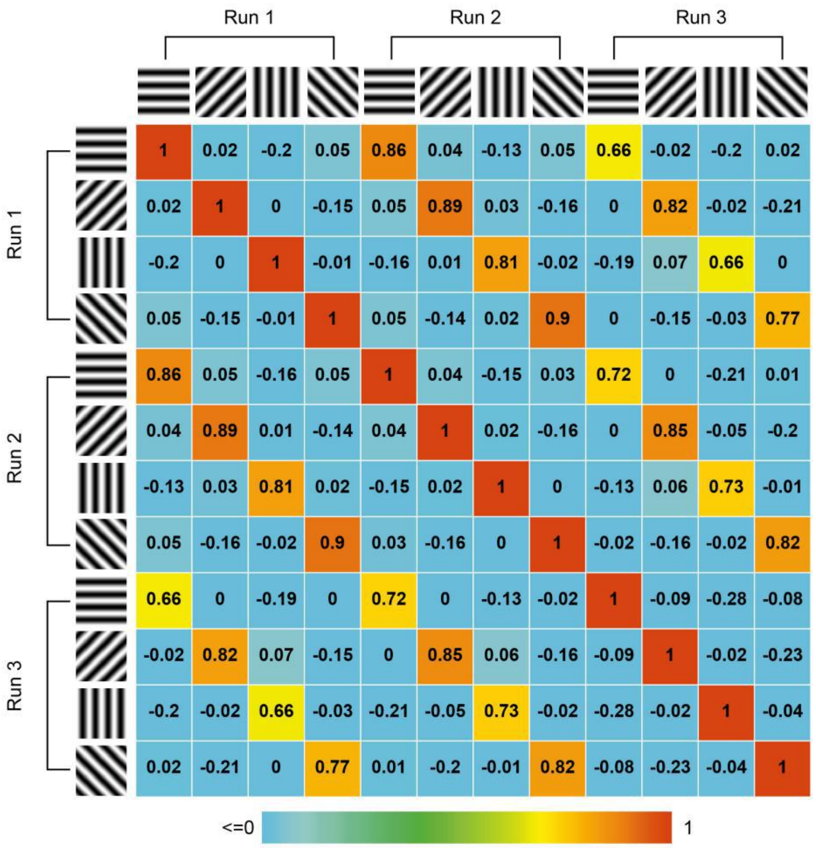


**Fig. S10. Correlation values between pairs of straight orientation maps obtained in 3 different runs (from Case 1).** These values are provided to give reader an idea of what high map correlations values are. When results from different runs are compared, only maps of similar orientation have high (orange, yellow) correlation values, while non-matching orientations produce low (blue) values. Color bar: correlation values (high, red, to low, blue) for pairs of maps. Note that negative values often occur for orthogonal orientations and are weaker in magnitude than positive values.

**Fig. S11.**

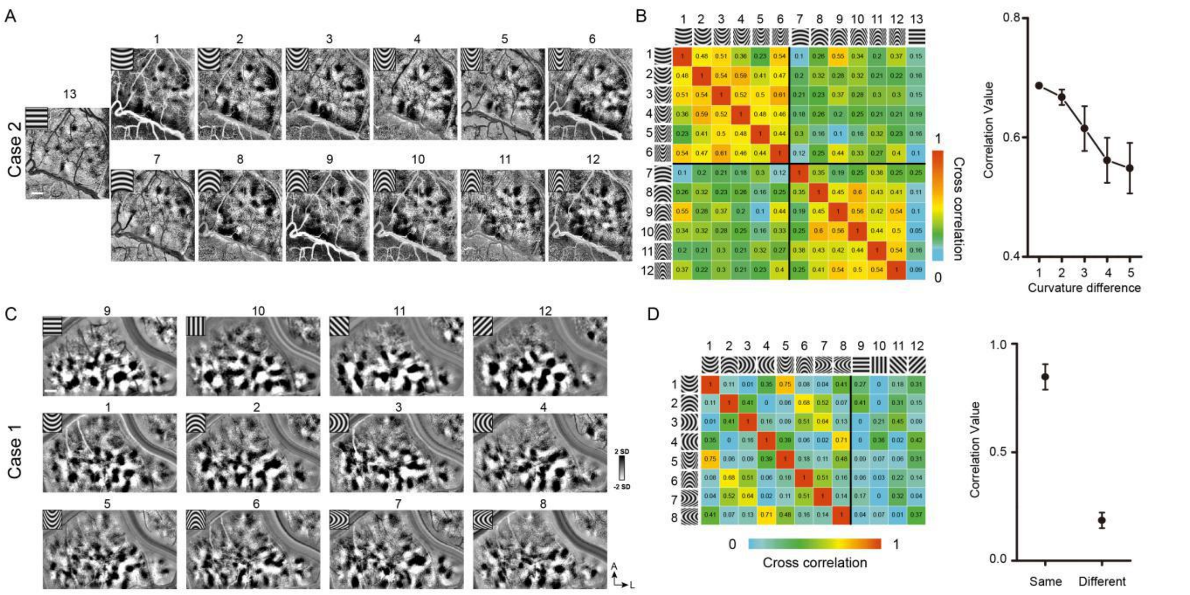


**<u>Fig. S11.</u> Systematic maps of curvature degree and curvature orientation in other cases.** A. Case 2: Maps of different curvature degrees. B. Correlation values for pairs of curvature response maps (from A). The more similar the curvature, the greater the correlation value (from B). C. Case 1: Imaging maps of different curvature orientations. D. Correlation values for pairs of curvature response maps (from C). Curvatures with the same orientation have greater correlation values (from D). Error bar: SEM. Scale bar: 1 mm.

**Fig. S12.**

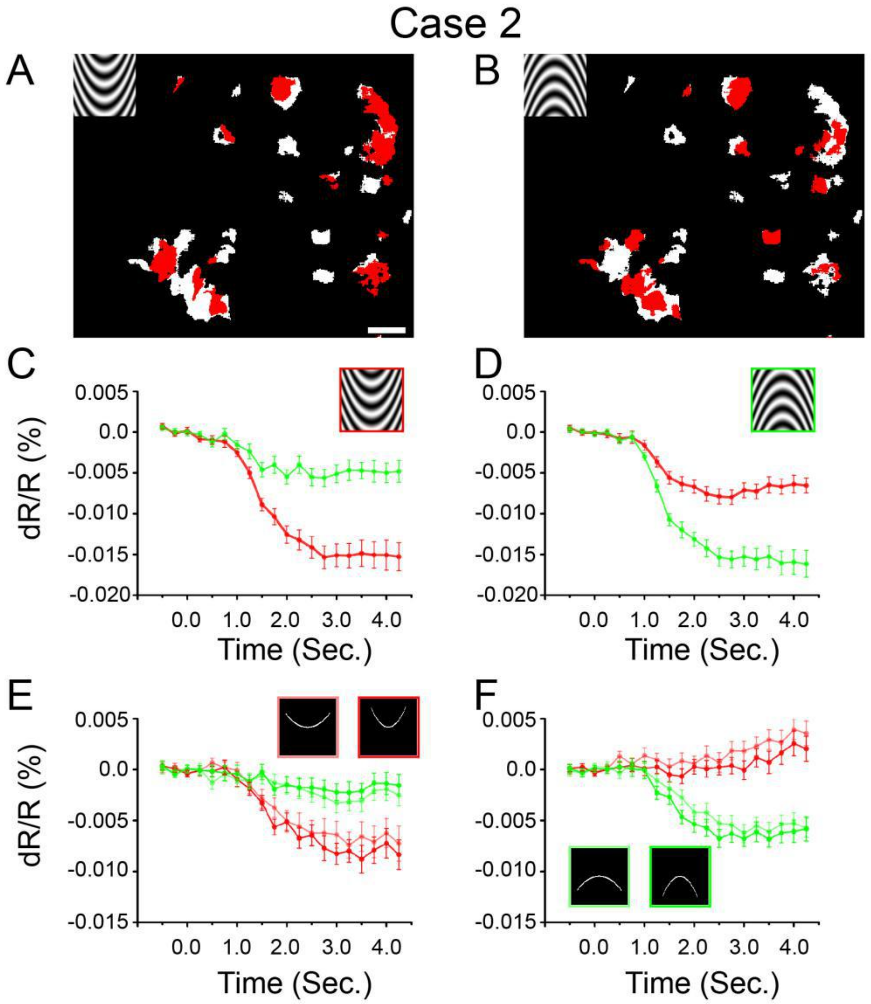


**<u>Fig. S12.</u> Curvature orientation cue-invariance from another case (Case 2).** A, B. Curvature orientation maps. C, D. Response timecourses of red pixels in maps above to curved gratings. E, F. Response timecourses of the same red pixels in maps above to curved single lines. Red lines: responses to up curvature, Green lines: responses to down curvatures. For flashed curved lines, we used two different curvature degrees. Scale bar: 1 mm.

**Fig. S13.**

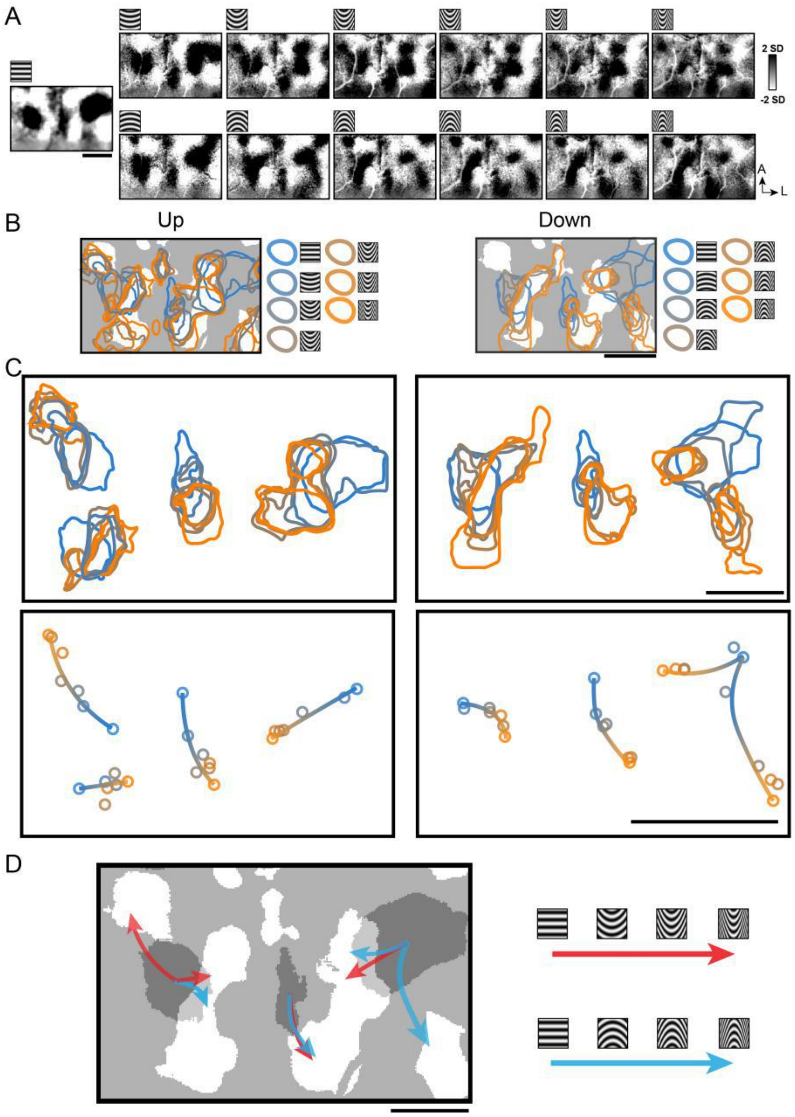


**<u>Fig. S13.</u> The activation shifts according to the curvature degree change.** A. Maps of different stimuli minus sum of straight gratings (Case 1). Top row: upwards curvatures, bottom row: downwards curvatures. Leftmost: horizontal gratings. B. Activated regions (two-tailed t-test, $p<0.01$) corresponding to different curvature stimuli. White regions: curvature domains. Color code: high (orange) to low (blue). C. Enlarged view of the activated region in B. Top row: activated regions (outlined by different colored lines). Bottom row: Shifting progressions of the activation center from straight orientation (blue dot) to low curvature (blue-orange dot) to high curvature (orange dot) are indicated by colored line (shaded from blue to orange). [Note that, the two leftmost domain progressions in Up panel are associated with the same orientation domain (blue) but are separated for clarity.] D. The centers of the activation shifted according to the change of curvature degree. White regions: curvature domains. Dark gray regions: horizontal orientation domains. Arrows: shifting directions of the activation from low to high (Red, upwards curvatures, Blue, downwards curvatures). Scale bar, 1mm.

**Fig. S14.**

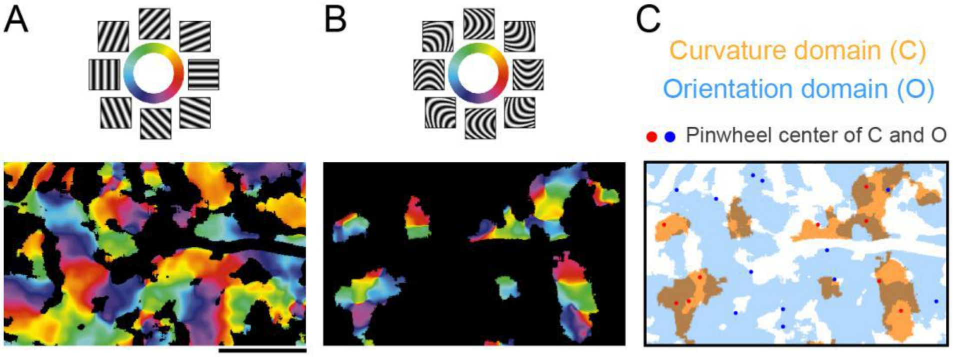


**<u>Fig. S14.</u> Curvature orientation and straight orientation maps from Case 3.** A. Polar map of straight grating response. Colored regions: orientation domains (significant response differences between two orthogonal orientations). B. Polar map of curved grating response. Colored regions: curvature domains (significant response differences between curvature gratings and straight gratings). C. Overlay of curvature response (orange) and straight orientation response (blue) zones. Red dots: curvature pinwheel centers, Blue dots: straight orientation pinwheel centers.

**<u>Table S1.</u> Case list**

| | Case 1 | Case 2 | Case 3 |
|---|---|---|---|
| Curvature vs. straight map | Fig. 1E | Fig. S5C,D | Fig. S6 |
| Curvature domain mask | FIg. 1G red pixels | Fig. 3A Right panel | Fig. 5A |
| Curvature degree map | Fig. 2 | Fig. S11A,B | NA |
| Curvature orientation map | Fig. S11C,D | Fig. 4 | Fig. 6, Fig. S14 |
| Cue (curvature orientation) invariance | NA | Fig. S12 | Fig. 5 |
| Scrambled response | NA | NA | Fig. S6 |

**Table S2.** **Statistic analysis of the response differences (related to Fig. S9).**

| | single curved line (Response AmplitudedR/R×$10^{-3}$ %) | single straight line (Response Amplitude dR/R×$10^{-3}$ %) | Wilcoxon test (curved vs. straight) P value |
|---|---|---|---|
| Case 2 | | | |
| Case 2 Curvature domain | -5.4 | -0.62 | $<10^{-5}$ |
| Case 2 Horizontal domain | 0.9 | -1.5 | <0.05 |
| Case 2 Veritical domain | 1.4 | -0.4 | >0.3 |
| Case 2 High spatial frequency domain | 4.0 | 1.4 | <0.05 |
| Case 3 | | | |
| Case 3 Curvature domain | -3.1 | 2.5 | $<10^{-5}$ |
| Case 3 Horizontal domain | 0.01 | -0.7 | <0.05 |
| Case 3, 45 deg orientation domain | -0.46 | -0.46 | >0.3 |
| Case 3, Veritical domain | 0.66 | 0.62 | >0.2 |
| Case 3, 135 orientation domain | 0.26 | -0.12 | >0.5 |
| Case 3 High spatial frequency domain | 0.68 | 0.54 | >0.3 |

**Table S3.** **Regression values for Fig. 2B.**

| | Condition | Length/width | Comparison pairs | Regression $R^2$ |
|---|---|---|---|---|
| Up | 1 | 1 | 2-1 3-1 4-1 5-1 6-1 | 0.68 |
| | 2 | 2 | 2-1 3-2 4-2 5-2 6-2 | 0.74 |
| | 3 | 4 | 3-1 3-2 4-3 5-3 6-3 | 0.74 |
| | 4 | 5 | 4-1 4-2 4-3 5-4 6-4 | 0.95 |
| | 5 | 7 | 5-1 5-2 5-3 5-4 6-5 | 0.92 |
| | 6 | 10 | 6-1 6-2 6-3 6-4 6-5 | 0.88 |
| Down | 7 | 1 | 8-7 9-7 10-7 11-7 12-7 | 0.9 |
| | 8 | 2 | 8-7 9-8 10-8 11-8 12-8 | 0.95 |
| | 9 | 4 | 9-7 9-8 10-9 11-9 12-9 | 0.82 |
| | 10 | 5 | 10-7 10-8 10-9 11-10 12-10 | 0.99 |
| | 11 | 7 | 11-7 11-8 11-9 11-10 12-11 | 0.96 |
| | 12 | 10 | 12-7 12-8 12-9 12-10 12-11 | 0.95 |
